# TelomereHunter: telomere content estimation and characterization from whole genome sequencing data

**DOI:** 10.1101/065532

**Authors:** Lars Feuerbach, Lina Sieverling, Katharina I. Deeg, Philip Ginsbach, Barbara Hutter, Ivo Buchhalter, Paul A. Northcott, Peter Lichter, Stefan M. Pfister, David T.W. Jones, Karsten Rippe, Benedikt Brors

**Author notes:** Equal contributors (Lars Feuerbach, Lina Sieverling).

## Abstract

**Summary:** Telomere shortening plays an important role in cellular aging and tumor suppression. The availability of large next-generation sequencing cohorts of matched tumor and control samples enables a computational high-throughput analysis of changes in telomere content and composition in cancer. Here we describe a novel software tool specifically tailored for the processing of large data collections.

**Availability and Implementation:** TelomereHunter is implemented as a python package. It is freely available online at:
http://www.dkfz.de/en/applied-bioinformatics/telomerehunter/telomerehunter.html.

## 1 Introduction

Telomeres are nucleoprotein complexes at the ends of eukaryotic chromosomes. In humans, telomeric DNA consists mainly of non-coding t-type (TTAGGG) repeats. However, c-(TCAGGG), g-(TGAGGG) and j-type (TTGGGG) variant repeats as well as other variations of the hexameric sequence exist (Coleman, et al., 1999; Lee, et al., 2014; Varley,et al., 2002). Telomeres shorten with each cell division and once a critical telomere length is reached, a DNA damage response is triggered, resulting in cellular senescence or apoptosis.

To circumvent the limited number of possible cell divisions, tumors employ activation of telomerase (Kim,et al., 1994) or “alternative lengthening of telomeres” (ALT) (Bryan,et al., 1997) as telomere maintenance mechanisms. Telomerase is an enzyme that adds t-type repeats to the chromosome ends. In contrast, ALT is based on recombination of telomeric regions and results in several characteristics, including telomeres of heterogeneous length and sequence composition.

Telomere maintenance mechanisms are crucial for tumorigenesis, making them valuable drug targets for cancer therapy (Shay, 2016). However, to precisely identify and interfere with these mechanisms in various tumor types, more insight into the different telomere structures is needed. In the last decades, several experimental methods have been established to assess telomere length and ALT status, e.g. terminal restriction fragment (TRF) analysis and telomere qPCR (Aubert, et al., 2012).

With the advance of massively parallel sequencing, an alternative method for measuring telomere content has emerged. Several studies have already shown that the number of short reads containing telomeric repeats can be used to estimate telomere content in whole genome sequencing (WGS) data, and that the results are comparable to those of established experimental methods (Conomos,et al., 2012; Ding, et al., 2014; Nersisyan and Arakelyan, 2015;Parker, et al., 2012).Here, we present TelomereHunter, a new computational tool for determining telomere content that is specifically designed for matched tumor and control pairs. In contrast to existing tools, TelomereHunter takes alignment information into account and reports the abundance of variant repeats in telomeric sequences.

## 2 Implementation

TelomereHunter is written as a python package and takes WGS BAM files of single samples or matched tumor and control pairs as input. Several parameters can be set by the user with the default settings and workflow being described in the following.

In the first step of TelomereHunter, telomere reads containing at least *n* non-consecutive repeats (t-, c-, g-or j-type) are extracted (Fig. 1A). *n* is calculated for each read depending on the read length with the following formula: n = floor(read length • 0.06). The criterion of searching for six non-consecutive repeats in 100 bp long reads has been proposed previously (Lee,et al., 2014) and was also found suitable for the data presented in the present study.

In the second step, the reads extracted are categorized depending on the alignment coordinates. If reads are properly paired, the mapping position of the mate is considered for the sorting. In short, reads mapping to intrachromosomal regions, i.e. all chromosome bands except the first or last band, are defined as intrachromosomal reads. The subtelomeric fraction comprises telomeric reads mapped to the first or last band of a chromosome. Telomeric reads from paired-end data are classified as junction-spanning if one mate maps to a first or last chromosome band and the other mate is unmapped. All remaining unmapped reads are categorized as intratelomeric.

The telomere content is calculated as the fraction of intratelomeric reads per million reads. To account for GC biases in sequencing data, TelomereHunter determines a GC-corrected telomere content: Instead of normalizing by the total number of reads in the sample, the intratelomeric reads are divided by the number of reads with a GC content between 48-52%, which is similar to that of the canonical t-type repeat and has been suggested for the normalization of telomeric reads (Ding,et al., 2014).

The output of TelomereHunter includes several diagrams visualizing the results (see Supplementary Fig. 1 for examples).

**Fig. 1.**

TelomereHunter workflow and validation. (A) Extraction and subsequent sorting of telomere reads with TelomereHunter. (B) Comparison of estimated telomere content tumor/control log2 ratios in pediatric brain tumor samples using experimental methods (TRF and qPCR) and computational analysis with TelomereHunter.

## 3 Results

### 3.1 Validation

For validation, TelomereHunter was compared to established experimental methods for telomere content measurement (see Supplementary Methods). The telomere content of nine pediatric brain tumor samples (six medulloblastoma and three glioblastoma samples) was determined computationally and was measured by telomere qPCR and TRF analysis. To demonstrate that TelomereHunter correctly determines the telomere content of both ALT-positive and ALT-negative samples, we included samples with different ALT status in the validation samples (as determined by TRF and C-circle assay, Supplementary Fig. 2).

The experimentally determined telomere content estimation was highly correlated with the TelomereHunter results for the individual tumor and control samples (r = 0.90 for qPCR and r = 0.65 for TRF). The correlation was further improved by GC correction of the computationally determined telomere content (r = 0.94 and 0.72; Supplementary Fig. 3). All methods consistently predicted telomere content gain or loss in the tumor sample compared to the control (Fig. 1B). The only exception was MB175, which can be explained by different amounts of DNA in the experimental setup (see Supplementary Methods).

### 3.2 Benchmark

Performance comparison of TelomereHunter with Motif Counter and TelSeq (Supplementary (Fig. 4 and Table 1) revealed that all computational results correlated better with qPCR than with the TRF analysis. TelomereHunter outperformed the other tools in the correlation to qPCR. The correlation of tumor/control log2 ratios to the TRF analysis was similar for all tools and the direct correlation to the TRF analysis was best using TelSeq.

**Table 1.**

Benchmark showing the correlation of computation and experimental telomere content estimation

## 4 Discussion and outlook

TelomereHunter reliably determines telomere content from WGS data. In contrast to existing tools, it takes mapping information into account and is able to search for a combination of the most common telomere repeat types. Moreover, TelomereHunter visualizes the results and, by default, gives a summary of telomere composition. We anticipate that the combination of telomere content determination and telomere repeat variant analysis from WGS data provided by TelomereHunter will prove to be valuable for identifying and characterizing telomere maintenance mechanisms in primary tumor samples.

## Acknowledgements

We thank the DKFZ Genomics and Proteomics Core Facility for provision of sequencing services. The authors would also like to thank Elke Pfaff (DKFZ) for her help in DNA sample selection.

### Funding

The work was funded within project CancerTelSys [grant number 01ZX1302 to K.R. and S.M.P.] in the e:Med program of the German Federal Ministry of Education and Research (BMBF) and supported by the German Federal Ministry of Education and Science in the program for medical genome research [grant number: 01KU1001A,-B,-C, and-D]. The work of P.G. and B.H. was supported by the intramural funding program of the German Cancer Research Center.

### Authors contributions

LF was responsible for the conception of the study. LF, LS and PG were involved in the design and writing of TelomereHunter. LS carried out the bioinformatical analyses. KID performed qPCR, TRF and C-circle assays. DTWJ and PAN coordinated sample acquisition. PL oversaw sequencing of the samples. BH and IB were responsible for preprocessing of the data. Experimental design and execution were overseen by SP, KR, DTWJ and BB. LF and LS wrote the manuscript with contributions by KID and DTWJ. KID, BH, DTWJ, PL and KR critically reviewed the manuscript. All authors read and approved the final manuscript.

### Conflict of Interest

none declared.

## 5 Supplementary Methods

### 5.1 Whole genome sequencing

The WGS datasets analyzed in this study were obtained from the PedBrain ICGC project. Matching tumor and control samples were collected according to ICGC guidelines. The DNA libraries were prepared using Illumina paired-end sample preparation protocols and sequencing was performed on Genome Analyzer IIx and Illumina HiSeq 2000 instruments as previously described (Jones,et al., 2012;Sturm,et al., 2012). Reads were aligned to the GRCh37 reference from 1000 Genomes project using bwa mem version 0.7.8 with the option-T 0.

### 5.2 Computational telomere content estimation using Motif Counter and TelSeq

In addition to TelomereHunter analysis, telomere content was determined using Motif Counter (http://sourceforge.net/projects/motifcounter/) (Conomos,et al., 2012) with the parameters-s-u-q 0 and TelSeq (Ding,et al., 2014) with default settings.

### 5.3 Telomere quantitative real-time PCR

Telomere qPCR was conducted essentially as described previously (Cawthon, 2002; O'Callaghan, et al., 2008). In short, 10 ng DNA, 1X LightCycler 480 SYBR Green I Master, 500 nM forward primer and 500 nM reverse primer were added per 10 μl reaction. The primer sequences were: telo fwd, 5'-CGGTTTGTTTGGGTTTGGGTTTGGGTTTGGGTTTGGGTT-3'; and telo rev, 5'-GGCTTGCCTTACCCTTACCCTTACCCTTACCCTTACCCT-3'; 36B4 fwd, 5'-AGCAAGTGGGAAGGTGTAATCC-3'; and 36B4 rev, 5'-CCCATTCTATCATCAACGGGTACAA-3'. Cycling conditions (for both telomere and 36B4 products) were 10 min at 95°C, followed by 40 cycles of 95°C for 15 s and 60°C for 1 min. A standard curve was used to determine relative quantities of telomere repeats (T) to those of the single copy gene (S, *36B4* gene, also known as *RPLP0*). The T/S ratio was calculated for each sample (tumor and control) separately. The log2 ratio of telomere content was determined by dividing the T/S ratio of the tumor sample by the T/S ratio of the control sample. The calculated log2 ratio represents the increase or decrease in telomere content in tumor versus control samples.

### 5.4 C-circle assay

The C-circle assay was performed according the protocol of Henson et al. (Henson,et al., 2009). Briefly, 30 ng DNA was combined with 10 μl 2X Φ29 Buffer, 7.5 U Φ29 DNA polymerase (both NEB), 0.2 mg/ml BSA, 0.1% (v/v) Tween 20, 1 mM each dATP, dGTP and dTTP and incubated at 30°C for 8 h followed by 20 min at 65°C. Reactions without addition of polymerase (-pol) were included as controls. After addition of 40 μl 2X SSC, the amplified DNA was dot-blotted onto a 2X-SSC-soaked Roti-Nylon plus membrane (Carl Roth). The membrane was baked for 20 min at 120°C and hybridized and developed using the TeloTAGGG Telomere Length Assay Kit (Roche). Chemiluminescent signals were detected using a ChemiDoc MP imaging system (Bio-Rad).

### 5.5 Terminal restriction fragment analysis

For TRF analysis, 4.5 μg genomic DNA of tumor and blood (control) samples were used, except for the GBM38 tumor and MB175 control sample, of which only 2.2 µg and 1.6 µg DNA were available, respectively. Genomic DNA was digested with the restriction enzymes HinfI and RsaI overnight. The digested DNA was resolved on a 0.6% agarose gel (Biozym Gold Agarose) in 1X TAE buffer using the CHEF-DRII pulsed-field gel electrophoresis system (Bio-Rad) with the following settings: 4 V/cm, initial switch time 1 s, final switch time 6 s, and 13 h duration. Southern blotting and chemiluminescent detection was performed using the TeloTAGGG Telomere Length Assay Kit (Roche) according to the manufacturer’s instructions. The blot was visualized with a ChemiDoc MP imaging system (Bio-Rad). The telomere content in each lane was determined by calculating the sum of intensities in each lane normalized to the amount of DNA loaded. This correction may not be sufficient if the difference of loaded DNA is too large. It is noted that qPCR and TRF differ with respect to the normalization between samples. For telomere qPCR, the telomere content is normalized to a single copy gene and thus has an internal control for the amount of DNA used. This control is lacking for the TRF analysis where only the total amount of DNA loaded is measured. Thus, the TRF analysis is more prone to errors that arise from differences in the amount of DNA between samples.

## supplementary figures

**Supplementary Figure 1.**
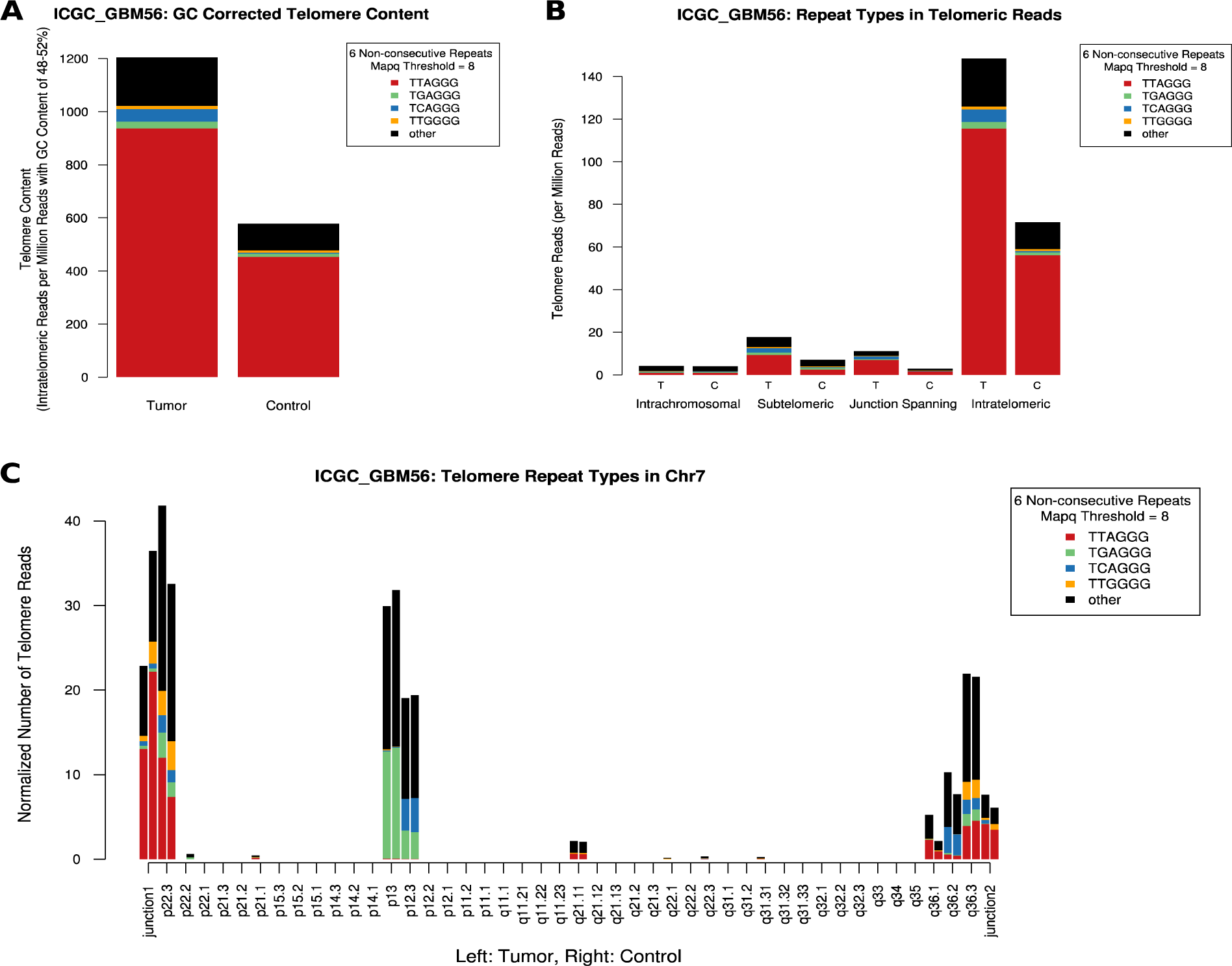
Exemplary output of TelomereHunter for a tumor sample and matching blood control of a glioblastoma patient ICGC_GBM56. (A) GC-corrected telomere content with amounts of different telomere repeat types, revealing that the c-type repeat is enriched in the tumor sample. (B) Number of telomere reads sorted into different fractions. (C) Junction-spanning and intrachromosomal reads mapped to chromosome 7 in ICGC_GBM56, indicating a pseudotelomeric region containing high numbers of g-and c-type repeats in bands p13 and p12.3. For chromosome bands, the number of telomere reads per million bases in the band and per billion reads in the sample is shown. Telomere reads assigned to junctions are shown per billion total reads. Similar plots are also made for other chromosomes. Stacks in the bar plots represent the relative occurrence of the searched repeat types.

**Supplementary Figure 2.**
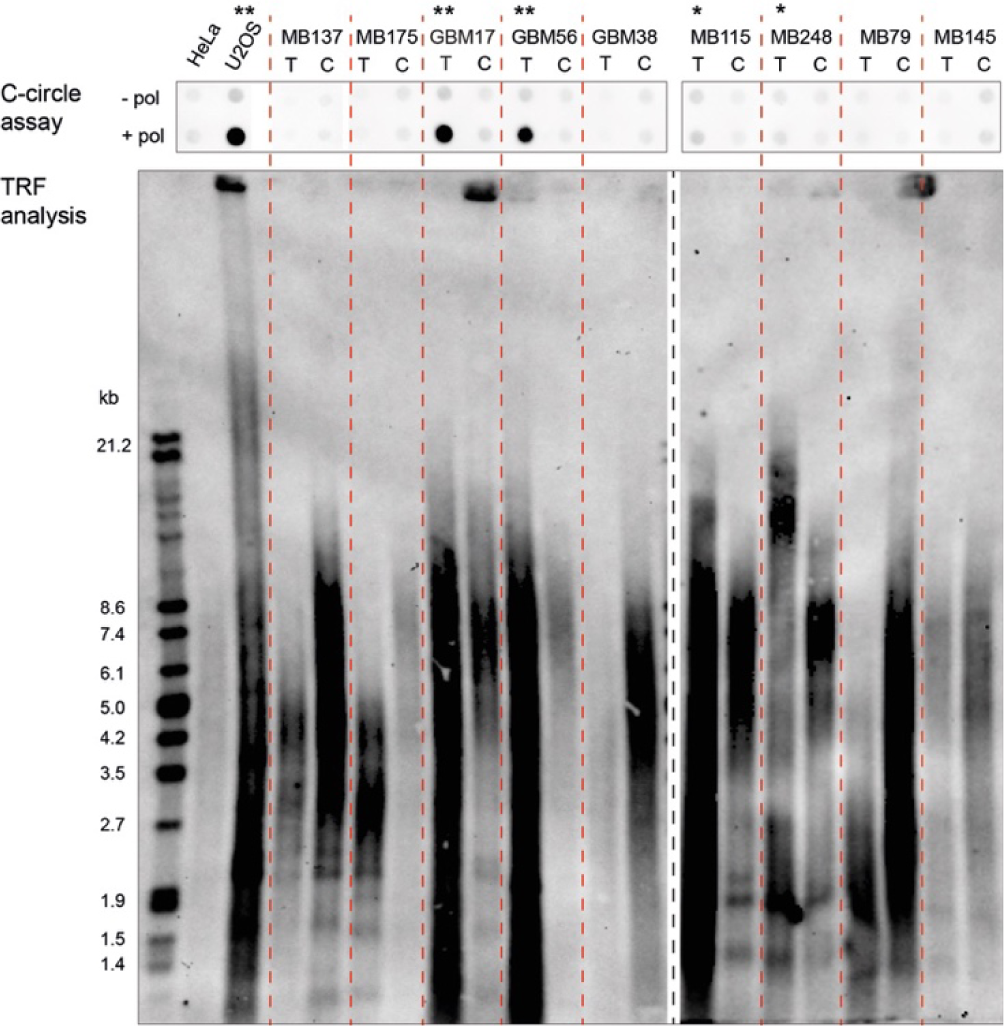
C-circle assay and TRF analysis of nine pediatric brain tumor samples (T) and matching controls (C). The ALT-negative HeLa and the ALT-positive U2OS cell line were included as a references. ALT-positive samples are highlighted by asterisks. * ALT-positive in TRF blot, ** ALT-positive in TRF blot and C-circle assay

**Supplementary Figure 3.**
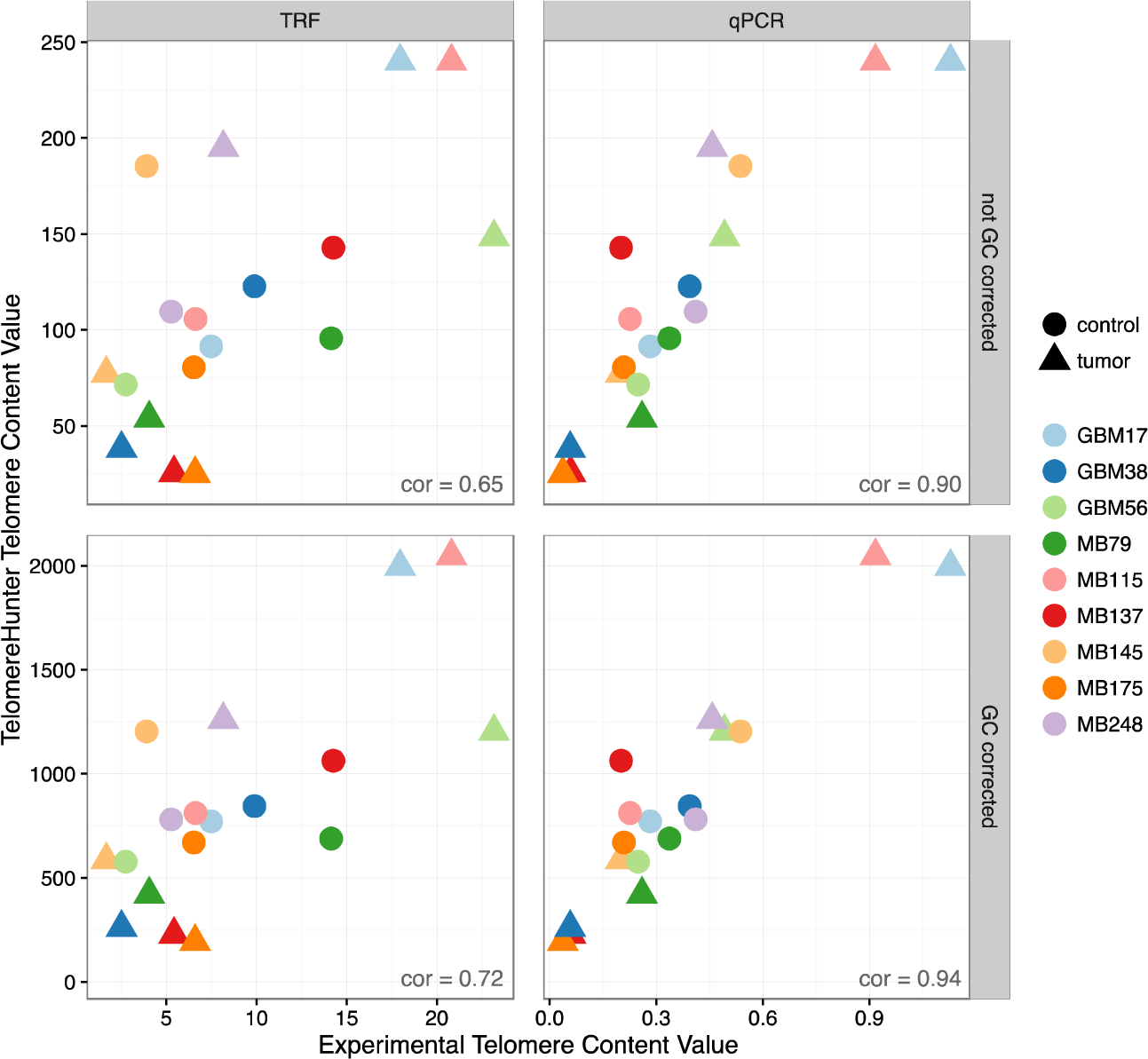
Comparison of estimated telomere contents in pediatric brain tumor samples and matching controls. The scatterplot shows the direct correlation between the telomere content estimated from WGS data with TelomereHunter and experimental telomere content estimation using TRF analysis and qPCR. The units of the GC-uncorrected and GC-corrected TelomereHunter results are intratelomeric reads per million total reads and intratelomeric reads per million reads with a GC content of 48-52%, respectively. Experimental telomere content values represent the summed intensities per *μ*g DNA for TRF analysis and the telomere to single copy gene (T/S) ratios for qPCR.

**Supplementary Figure 4.**
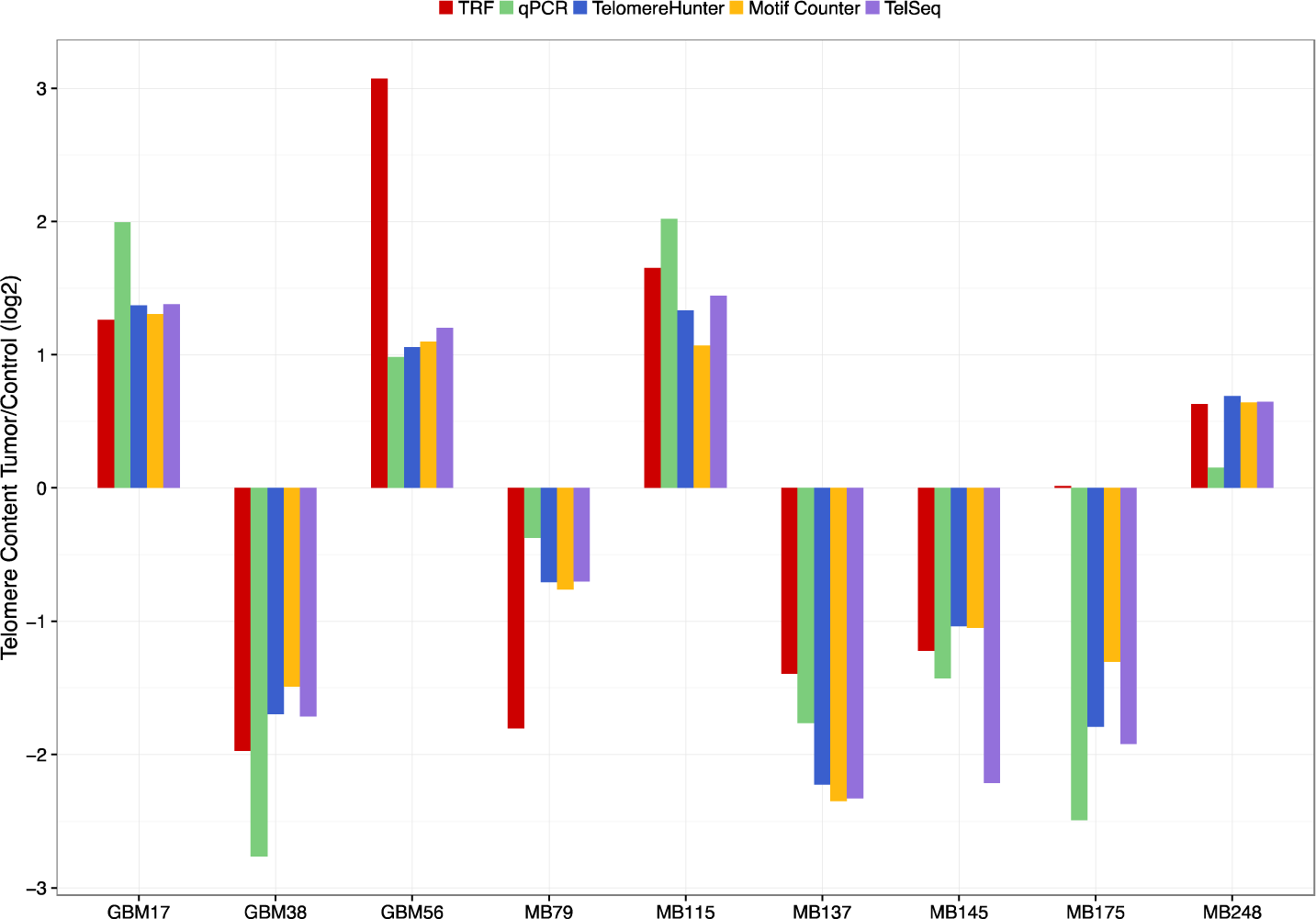
Comparison of experimental and computational telomere content estimation. Telomere content tumor/control log2 ratios were estimated for nine sample pairs from pediatric brain tumor patients using TRF, qPCR and the computational tools TelomereHunter, Motif Counter and TelSeq.

## References

Aubert, G., Hills, M. and Lansdorp, P.M. Telomere length measurement-caveats and a critical assessment of the available technologies and tools. Mutat Res 2012;730(1–2):59–67.

Bryan, T.M., et al. Evidence for an alternative mechanism for maintaining telomere length in human tumors and tumor-derived cell lines. Nat Med 1997;3(11): 1271–1274.

Coleman, J., Baird, D.M. and Royle, N.J.The plasticity of human telomeres demonstrated by a hypervariable telomere repeat array that is located on some copies of 16p and 16q. Hum Mol Genet 1999;8(9): 1637–1646.

Conomos, D.,et al.Variant repeats are interspersed throughout the telomeres and recruit nuclear receptors in ALT cells.J Cell Biol 2012;199(6): 893–906.

Ding, Z., et al. Estimating telomere length from whole genome sequence data. Nucleic Acids Res 2014;42(9):e75.

Kim, N.W., et al. Specific association of human telomerase activity with immortal cells and cancer. Science 1994;266(5193): 2011–2015.

Lee, M., et al. Telomere extension by telomerase and ALT generates variant repeats by mechanistically distinct processes. Nucleic Acids Res 2014;42(3): 1733–1746.

Nersisyan, L. and Arakelyan, A. Computel: computation of mean telomere length from whole-genome next-generation sequencing data. PLoS One 2015;10(4):e0125201.

Parker, M., et al. Assessing telomeric DNA content in pediatric cancers using whole-genome sequencing data. Genome Biol 2012;13(12):R113.

Shay, J.W.Role of Telomeres and Telomerase in Aging and Cancer. Cancer Discov 2016;6(6): 584–593.

Varley, H., et al. Molecular characterization of inter-telomere and intra-telomere mutations in human ALT cells. Nat Genet 2002;30(3): 301–305.

## References Supplementary Material

Cawthon, R.M. Telomere measurement by quantitative PCR. Nucleic Acids Res 2002;30(10):e47.

Conomos, D., et al. Variant repeats are interspersed throughout the telomeres and recruit nuclear receptors in ALT cells. J Cell Biol 2012;199(6):893–906.

Ding, Z.,et al.Estimating telomere length from whole genome sequence data. Nucleic Acids Res 2014;42(9):e75.

Henson,J.D., et al. DNA C-circles are specific and quantifiable markers of alternative-lengthening-of-telomeres activity. Nat Biotechnol 2009;27(12):1181–1185.

Jones, D.T., et al.Dissecting the genomic complexity underlying medulloblastoma. Nature 2012;488(7409):100–105

O'Callaghan, N.,et al. A quantitative real-time PCR method for absolute telomere length. Biotechniques 2008;44(6):807–809.

Sturm, D.,et al. Hotspot mutations in H3F3A and IDH1 define distinct epigenetic and biological subgroups of glioblastoma. Cancer Cell 2012;22(4):425–437.

